# Cascading effects of elephant-human interactions in a savanna ecosystem and the implications for ecology and conservation

**DOI:** 10.1101/2021.08.18.456886

**Authors:** David Western, Victor N. Mose

## Abstract

Our study monitored the changes in elephant numbers, distribution and ecological impact over a fifty-year period as the free-ranging intermingled movements of wildlife and traditional subsistence pastoralists across the Amboseli ecosystem were disrupted by a national park, livestock ranches, farms, settlements and changing lifestyles and economies.

Elephants compressed into the national park by poaching and settlement turned woodlands to grassland and shrublands and swamps into short grazing lawns, causing a reduction of plant and herbivore diversity and resilience to extreme events. The results echo the ecological findings of high-density elephant populations in protected areas across eastern and southern Africa. The impact has led to the view of elephants in parks being incompatible with biodiversity and to population control measures.

In contrast to Amboseli National Park, we found woody vegetation grew and plant diversity fell in areas abandoned by elephants. We therefore used naturalist and exclosure experiments to determine the density-dependent response of vegetation to elephants. We found plant richness to peak at the park boundary where elephants and livestock jostled spatially, setting up a creative browsing-grazing tension and a patchwork of habitats explaining the plant richness.

A review of prehistorical and historical literature lends support to the Amboseli findings that elephants and people, the two dominant keystone species in the savannas, have been intimately entangled prior to the global ivory trade and colonialism. The findings point to the inadequacy of parks for conserving mega herbivores and as ecological baselines.

The Amboseli study underlines the significance of space and mobility in expressing the keystone role of elephants, and to community-based conservation as a way to win space and mobility for elephants, alleviate the ecological disruption of compressed populations and minimize population management.

## Introduction

After decades of physical forces being viewed as bottom-up drivers of ecosystem processes (1–3) the role carnivores and herbivores in governing community dynamics from the top-down has also been well established in both ecological theory (4–8) and conservation policy and management (9–11). The extermination, reintroduction, displacement and compression of keystone large herbivores and carnivores can all have long-term repercussions on ecosystems and landscapes (12, 13)

Given the growing impact humans have on wildlife (14), population disruptions offer naturalistic experiments for testing the role of megamammals as keystone species precipitating trophic cascade through ecosystems (7). The impact of megafaunal disruptions will, however, differ between biogeographic regions depending on their histories. The heavily depleted faunal assemblages of North America during the Pleistocene (15) and Madagascar during the Holocene (16), for instance, will respond differently to East Africa’s megafauna which has remained relatively intact since the late Pleistocene (10, 17). Whereas the reintroduction of the wolf into Yellowstone’s Pleistocene-depleted predator fauna caused large ripple effects on the plant and animal community (11), the recolonization of wild dogs on the Laikipia Plateau in Kenya (18) in contrast, caused little discernable impact due to the present of other large predators, including lions and hyenas. How much greater then would be impact of reintroducing megaherbivores such as elephants and rhinos into North America biomes after 11,000 years of faunal depletion and weakened plant herbivore defenses than the reintroduction of the wolf into Yellowstone?

Africa offers an indication of the far greater biotic impact of large herbivores than carnivores on ecosystems. The ecological impact of the continent’s suite of megaherbivores on biomes ranging from forests to savanna has been well-documented (19–22). Most studies have, however, been conducted in protected areas where the impact of elephants especially has been intensified by human activity causing compressed populations and curtailed seasonal movements. Wall et al. (2021) showed the distribution of elephants to occupy only 17% of their potential range due to the level of the human disturbance and safety in protected areas. Normalized for rainfall, censuses of 34 populations showed elephant densities in East Africa to be five-times higher in protected than non-protected areas (24).

Discussions on the keystone role of elephants have centered on their impact on woody vegetation and whether the findings support equilibrium, non-equilibrium or alternative multi-state theories of ecosystem dynamics (25–27). Here again, using protected areas as templates exaggerates the deleterious impact of elephants on biodiversity and obscures the keystone role they play in free-ranging interactions with humans. The lack of such studies prior to the establishment of protected areas masks the complex and shifting interplay which likely shaped African ecosystems for millennia (28). Prehistorical and historical reviews of the interlinked ecologies help reveal the role elephants play in ecosystems when free-ranging rather than in compressed and fragmented populations.

## Elephant-human interactions

Paleo records and historical accounts point to the ancient links and changing relationship between hominins and elephants since the Lower and Middle Stone Age. The earliest evidence of human interactions with elephants comes from the fossil bones in Olorgesailie formation 150 km east of Amboseli in the Rift Valley. Here at 1 million BP, the fossilized remains of *Elephas recki*, a species twice the weight of *Loxodonta africana*, is surrounded by discarded stone tools and shows multiple cut marks on the bones (29). It is unclear if the disappearance of *Elephas recki,* which coincided with the 75% turnover of the large mammal fauna in the transition from the Lower to Middle Stone Age 500,000 to 350,000 BP, was due to climatic changes or the emergence of more sophisticated weaponry and hunting techniques or both (30). The eruption of taller grasslands following the loss of the largest species across a range of herbivore taxa at a time the climate was drying is, however, consistent with Owen-Smith’s (1987) herbivore release hypothesis explaining the growth of coarse vegetation following the extinction of megaherbivores.

The presence of sophisticated spears found among wild horse fossils preserved in a peat bog in Germany at 350,000 BP (32) adds further evidence of projectile weapons and group hunting as plausible explanations for the Olorgesailie faunal turnover. The transition marks the emergence of *Homo sapiens, Loxodonta africana* and the modern large mammal fauna (29) in which the keystone roles of elephants and humans have been linked over hundreds of millennia.

The Venus Hohle Fels figurines dating from 35,000 to 40,000 BP testify to the role of mammoth ivory in prehistoric cultural practices following the arrival of modern sapiens in Europe (33) and most likely among Neanderthals far earlier. Historical accounts document a flourishing North African ivory trade supplying the Greeks and Romans from as early as 2,500 BP, and the extirpation of elephants from their north African ranges during the classical times (34). The dhow trade driven by the Indian Ocean trade winds transported large quantities of ivory from Africa to the Arabian Peninsula and India from 500 AD onwards, reaching a peak in the mid-19^th^ century using slaves as portage (35–37). Evidence points to the depletion of elephant populations across eastern and southern African by the late 19th century (37, 38), resulting in the growth of woody vegetation (39). Alarmed by the outlook for elephants, the eight colonial powers led by the British and German governments convened the first ever wildlife conference in 1900 to save wildlife, and elephants particularly, from the extinctions happening in southern Africa (40).

Historical records of Amboseli mirror the continental picture. The writings of Joseph Thomson and Count von Hoehnel, the first European explorers to pass through Amboseli in the 1880s, make no mention of seeing elephants (41, 42). Photographs by Schillings (1906) in the early 1900s show an abundance of regenerating fever trees (*Acacia xanthophloea*) but no mature woodlands. The elephant population rose steadily from a low point at the turn of the 20^th^ century in response to the abolition of the slave trade and conservation protection by the British colonial administration, to a peak in the late 1960s (37, 44). A repeat photographic survey of photos taken by Martin and Osa Johnson in 1921 (45), shows the thick regenerating groves of fever trees and a dearth of elephant dung or tree damage. In the years following the trees matured and elephant damage intensified, leading to the replacement of woodlands by grasslands across much of the Amboseli Basin from the 1950s onwards (46).

Gazetted as Amboseli National Reserve in 1948, the Amboseli Basin at the center of the reserve was renowned for its majestic fever trees framing the backdrop of Kilimanjaro (47). The film, *Where No Vultures Fly* (48), shot extensively in Amboseli in 1950, showed growing signs of elephant damage to the fever tree groves. Early wardens reported a rising number of elephants in the Amboseli Basin that period (47). By the early 1960s conservationists began voicing concerns over the rapid loss of the iconic yellow fever trees and pressured government to declare Amboseli a national park to protect the woodlands in the belief that overstocking by Maasai livestock was the cause (49). Later studies suggested other causes, including elephant damage (19), rising salinity accelerated by elephant damage (50), synchronized woodland growth and ageing (51) and hydrological causes (52). Long-term, experimental studies (53) and ecological monitoring (54) factored out elephants as the main cause of woodland loss and broader habitat changes.

Given that elephants have contracted to a fifth of available habitat due human displacement (23), how did the interaction of elephants and humans play out before the industrial and colonial age (28), how has their interaction change as a result, and how can the space and mobility needed to avoid compression and ecological degradation be reestablished?

The Amboseli Conservation Program (ACP) offers some insights on these questions. The monitoring and research program began in 1967 when Maasai pastoralists still practiced subsistence husbandry and moved seasonally across communally owned land. The early study period preceded the division of the ecosystem into a national park and group ranches (49, 55). Here we draw on the half century of monitoring and research to document the push and pull forces of human agencies on the movements, residency times, densities, foraging patterns and complex ecological cascades they triggered by elephants.

## Methods

### Study area

The 3,700 km^2^ part of the Amboseli ecosystem (Fig 1) is defined by the seasonal movements of the large wild herbivore and pastoral livestock populations lying in the rain shadow of Kilimanjaro along the Kenya-Tanzania border. The wild herbivore migrants, including elephants, zebra, wildebeest, hartebeest, eland and buffalo, move seasonally between the wet season and the dry season range concentrated around the permanent swamps of the Amboseli Basin. The ecosystem has been detailed in several publications (56–59). The vegetation is dominated by bushed grassland falling within ecological Zone V of Pratt, Greenway and Gwynne (60). Aquifers emanating in the northern forests of Kilimanjaro drain into the dry Pleistocene lakebed of the Amboseli Basin, creating a series of permanent swamps and a shallow water-table which supports hydrophilic vegetation dominated by *Acacia xanthophloea* woodlands. Maximum temperatures range from 26 to 44°C and minimum temperatures from 6 to 14°C. The twice-yearly rain seasons fall between October to December and March to May, averaging 350mm annually (61).

**Fig 1:**
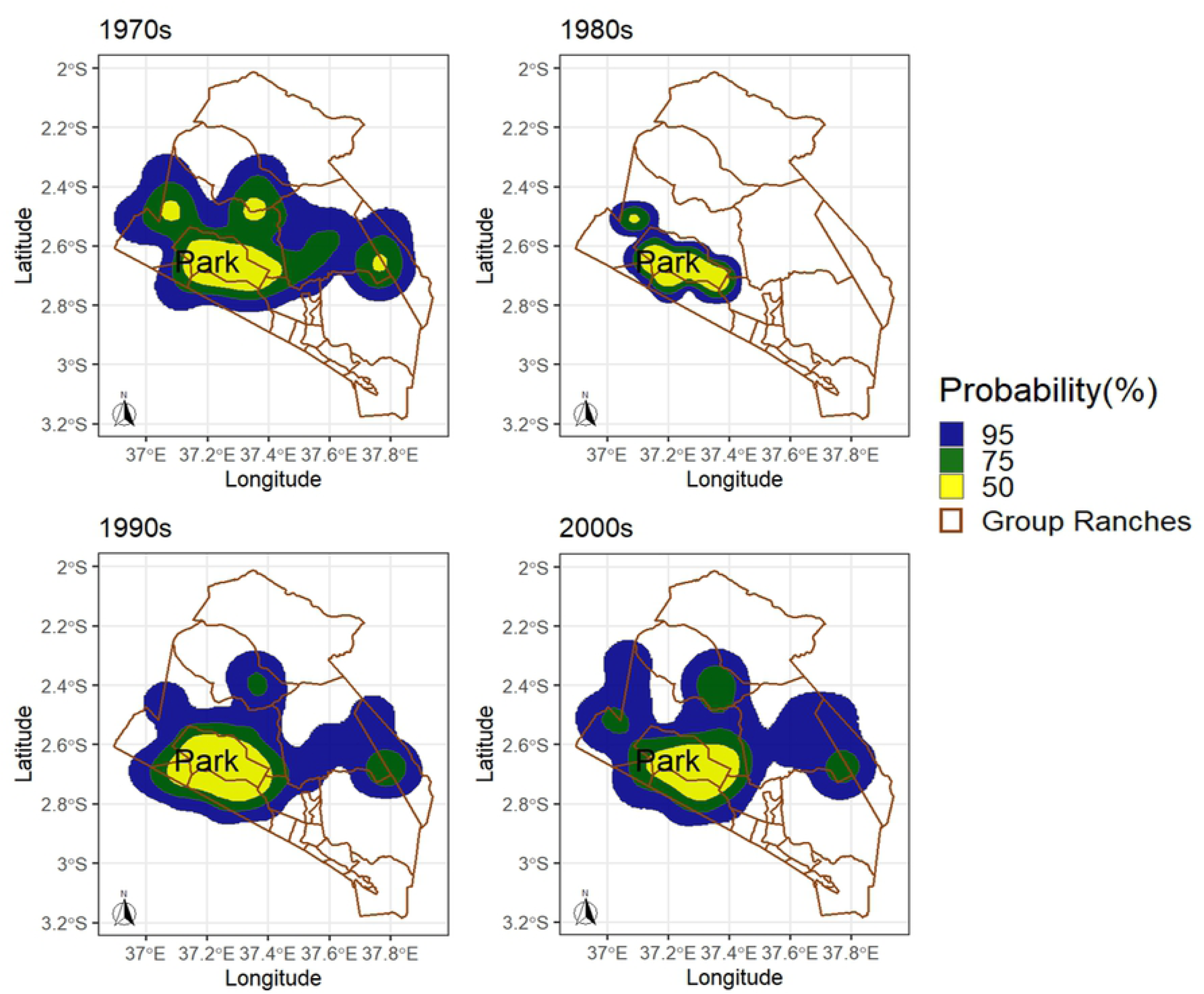
Kernel Density Distributions of Elephants in Amboseli Ecosystem, Kenya 1974 to 2016.

Despite the abundance of wildlife, the herbivore biomass is dominated by the cattle, sheep and goats of the pastoral Maasai who followed the same seasonal migratory patterns as wildlife until the late 1970s. In 1977 livestock were excluded from the 388 km^2^ Amboseli National Park to protect the rich wildlife concentrations of the Amboseli Basin and secure their late season forage against settlement and farming. The Maasai in the surrounding communal lands were given title deeds to seven group ranches, each managed separately by elected representatives (55). The higher rainfall slopes of Kilimanjaro were divided into small farms which spread downslope towards the permanent swamps from the 1960s onwards. Beginning in the 1970s the permanent swamps Namelog and Kimana lying east of Amboseli National Park were subdivided into private holdings supporting irrigated farming. Subdivision of the group ranches, which began in the northern Kaputei Section of eastern Kajiado District in the 1970s and displaced wildlife including elephants (55), is currently underway in the group ranches surrounding Amboseli National Park.

### Herbivore monitoring

Sample aerial and ground counts of the 600 km^2^ dry season concentration area of the Amboseli basin were conducted between 1967 and 1971 using the methods described in Western (1973) and Pennycuick and Western (1972). Total aerial counts of elephants, buffalo and human settlements in the basin area have been flown from 1975 onwards using methods described by Western and Lindsay (1984). Aerial sample counts of the 8,500 km^2^ eastern Kajiado county covering the Amboseli ecosystem and bordering areas in eastern Kajiado County, have been flown one to several times most years from 1974 and 2020. The counting methods have been described in detail elsewhere (64–67). The area was divided on a UTM map projection into 5 by 5 km grids. Flight lines were flown through the center of each grid in a north south-direction 90m above ground with two back-seat observers counting all wild and domestic herbivores 25kg upwards, as well as settlement structures within 150 to 200 meters. Herds too large to count by eye were photographed and later counted under a binocular microscope. Population estimates were derived from the 8% to 10% sample counts using the Jolly II equation (68). We used Douglas-Hamilton and Burrill’s (1991) ratio of dead to live elephants to measure poaching levels. The long-term changes in large herbivore populations from 1974 to 2020 have been detailed in Western and Mose (2021).

Human activity, including settlements, the number and type of huts and presence of farming in a grid, was recorded for each grid square. Traditional Maasai dung huts are a useful measure of seasonally mobile settlements, and thatch and tin huts and houses of permanent settlements. We use settlement type, numbers and proportion of a grid covered by farms as measure of human activity rather than human population size, given that livestock and farms most directly influence the numbers and distributions of wildlife (65, 69).

### Vegetation monitoring

The detection of woodland change in the Amboseli Basin was drawn from data measuring tree density in 30 randomly distributed 18ha plots demarcated on aerial photos dated 1950, 1961, 1967 and 1980. Tree condition was measured in 18 plots using the ratio of dead to live limbs and elephant damage based on the proportion of stripped bark. The methods were described by Western and Van Praet (1973). A further count of tree density was conducted along the two transects using 2020 Google Earth satellite imagery. The density of elephants along the transects were averaged from monthly total counts of the Amboseli Basin between 1975 to 1988 using 1km^2^ grid. Further woodlands density and associated vegetation composition studies were conducted in 1988 along transects running east and west at 0.75km intervals from the center of the Amboseli Basin. Tree cover and bush density by species was measured by the point-center-quarter method (70) and herb layer composition by the slanting pin frame method (71). More detailed descriptions of the vegetation structure, composition and long-term changes can be found in Western et al. (2021).

## Results

### Changes in distribution and population size

The decadal changes in elephant distribution from the 1970s to 2000s are shown in Fig 2. The distribution maps confirm the earlier findings of Western and Lindsay (1984) depicting two overlapping populations, one centered on Amboseli Basin, the other on Tsavo West and Chyulus Hills National Parks. In the 1980s the two populations contracted sharply and by the 1990s had split into two discrete populations concentrated in Amboseli and Tsavo West National Park. By the 2000s both populations had substantially recovered their 1970s ranges and overlapped once again east of Amboseli. The annual range of the Amboseli sub-population based on the kernel density maps declined from 7159 km^2^ in the 1970s to 1984 km^2^ in the 1980s, a 72% shrinkage. The range subsequently expanded to 5916 km^2^ in the 1990s and 6,349 km^2^ in the 2000. Satellite collared animals show elephants ranging into Tanzania and across to the Rift Valley (73) by 2018, a far more extensive range than elephants radio tracked in the 1970s (63).

**Fig 2:**
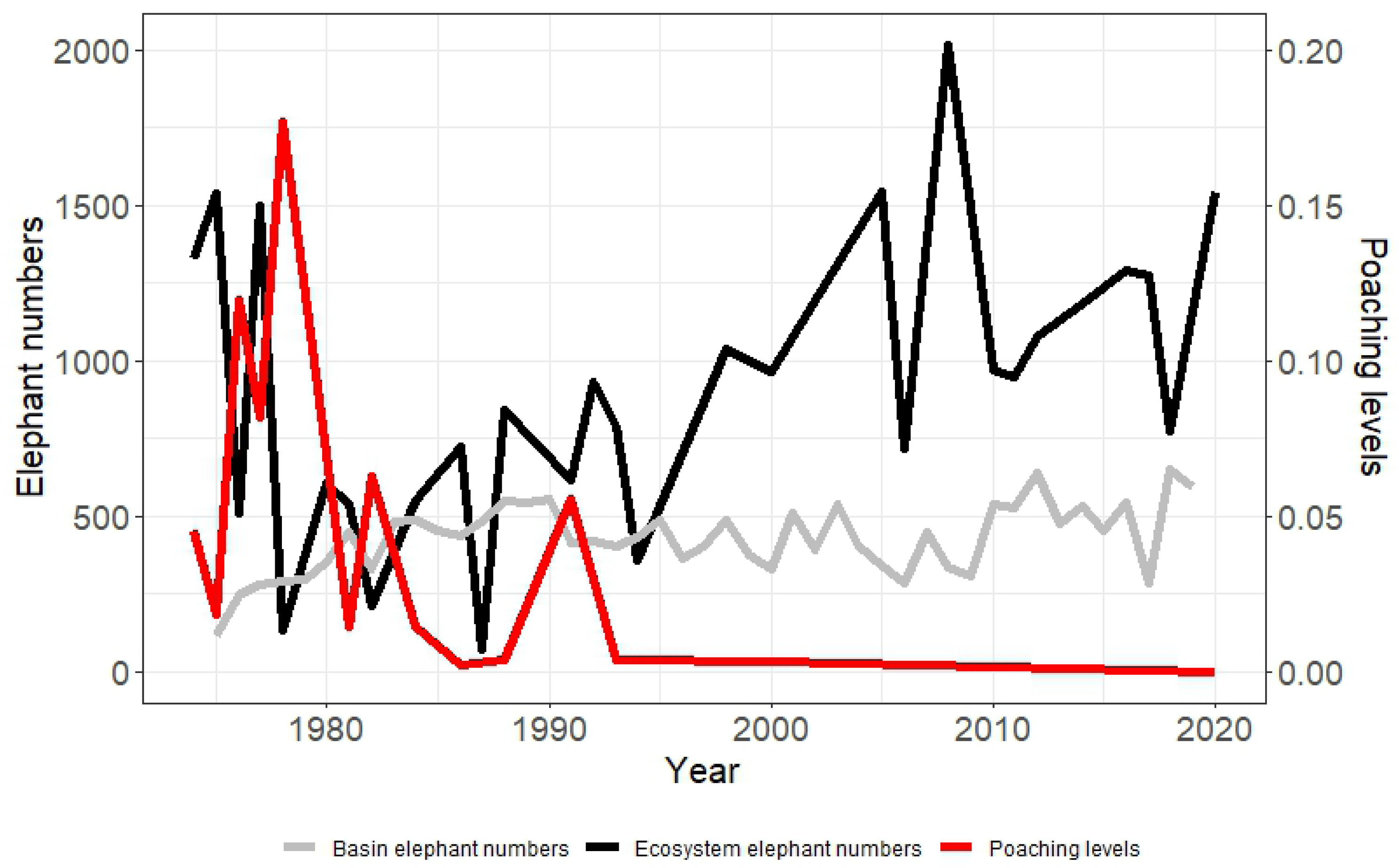
Elephant numbers in the 8,500 km^2^ of eastern Kajiado spanning the Amboseli ecosystem (black) and 700 km^2^ dry season range (grey) plotted against carcass ratios as a measure of poaching levels (red).

Fig 2 shows the elephant numbers and carcass ratios in the 8,500km^2^ ecosystem based on sample aerial counts from 1974 to 2020. The numbers in the 600km^2^ dry season range are based on sample ground counts and aerial counts from 1968 to 1971, and total aerial counts from 1975 onwards. The counts are averaged for each year where more than one count was conducted.

Elephant numbers in the ecosystem fell from between 1,200 and 1,500 in the early 1970s to under 500 between 1974 and 1978 (Fig 2). We attribute the precipitous loss to poaching for several reasons. First, at the onset of the aerial counts of the ecosystem, large numbers of recently dead elephants with their tusks hacked out were found across the ecosystem outside Amboseli National Park. Carcass counts were added to the live animal counts to document the numbers, distribution and approximate age of dead elephants. Second, the population decline is inversely related to the poaching rate measured by the carcass: live ratio (74) over the previous one year (*r* = −0.37, *P* = 0.0453). Third, the peak number of elephant carcasses recorded from the aerial counts at the height of poaching in 1976 numbered 536±102. Allowing for many undetected carcasses due to decomposition and scattering by predators over the previous three years, and others hidden under dense bush, the number of poached elephants explains most of the population losses. Fourth, the recovery of the elephant population coincides with the decline in poaching (Fig 2). Finally, the timing of the poaching surge matches the wholesale slaughter of elephants throughout Kenya and eastern and central Africa following a surging increase in ivory prices on the world market in the early 1970s (74, 75).

The distribution of carcasses during the poaching surge of the 1970s (Fig 3) depicts the most vulnerable locations: in the west in the dense bush country stretching to Namanga, in the north and northeast on the fringes of the elephant range, and in the east on the Chyulu Hills and southeast on Rombo Group Ranch, both areas where elephants spread out from Tsavo East National Park. A small number of elephants were poached south of the park boundary and the few carcasses were recorded in the park, largely young animals which succumbed to the mid-1970s drought (54). In contrast to the widespread abundant carcasses recorded in the 1970s. the number dropped to a few incidental records in the 2000s (Fig 2).

**Fig 3:**
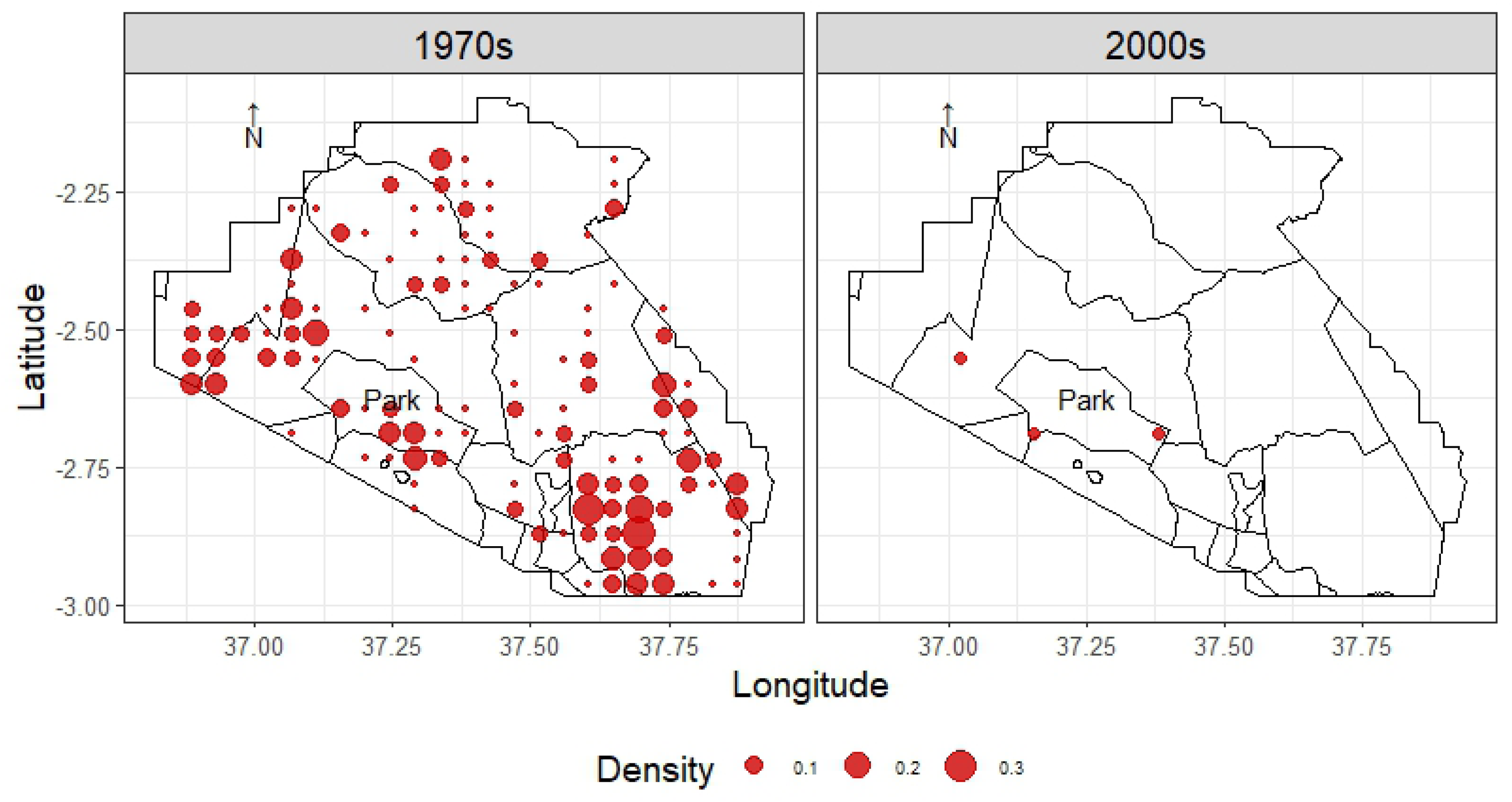
The density distribution of elephant carcasses in the 1970s at the height of poaching and after the 2000s when elephants recolonized their former range (Fig 1).

We attribute the rising numbers of elephants in the park through the 1970s and early 1980s (Fig 2) to elephants fleeing poachers to the safety of the national park (59) where the concentration of tourist vehicles and ranger forces offered protection. The safety factor was also evident in the relaxed behavior of herds in the park and their clumped and agitated formations in moving out of the park (63) and in response to human disturbances (76). Most movements out of the park were nocturnal, a pattern common to other protected area populations in response to human threats (23).

Another factor which may have added to the safety of the park was the exclusion of livestock after the creation of Amboseli National Park. Though formally gazetted in 1974, livestock were not banned from the park until 1978 when alternative water sources were made available (77). By then, elephant numbers had been rising rapidly since the early 1970s (Fig 4). Although the Maasai moved back into the park in 1983 after the government failed to honor an annual payment from park fees (78), elephant numbers continued to rise through to the 1990s when the seasonal migrations resumed though in reverse. Elephants migrated into the park during the rains and left in growing numbers with the advance of the dry season. The dry season exodus continued through the 1990s and into the 2000s (Fig 4). The growing population and resumption of migrations saw elephants recolonized their former range (Fig 4) due largely to a drop in poaching following the deployment of community rangers across the surrounding group ranches. The reversal in the wet season migrations typifying elephants, zebra, wildebeest and livestock in the 1960s and 1970s point to explanations other than poaching. We explore alternative explanations after looking at the habitat changes caused by elephant compression into the park.

**Fig 4:**
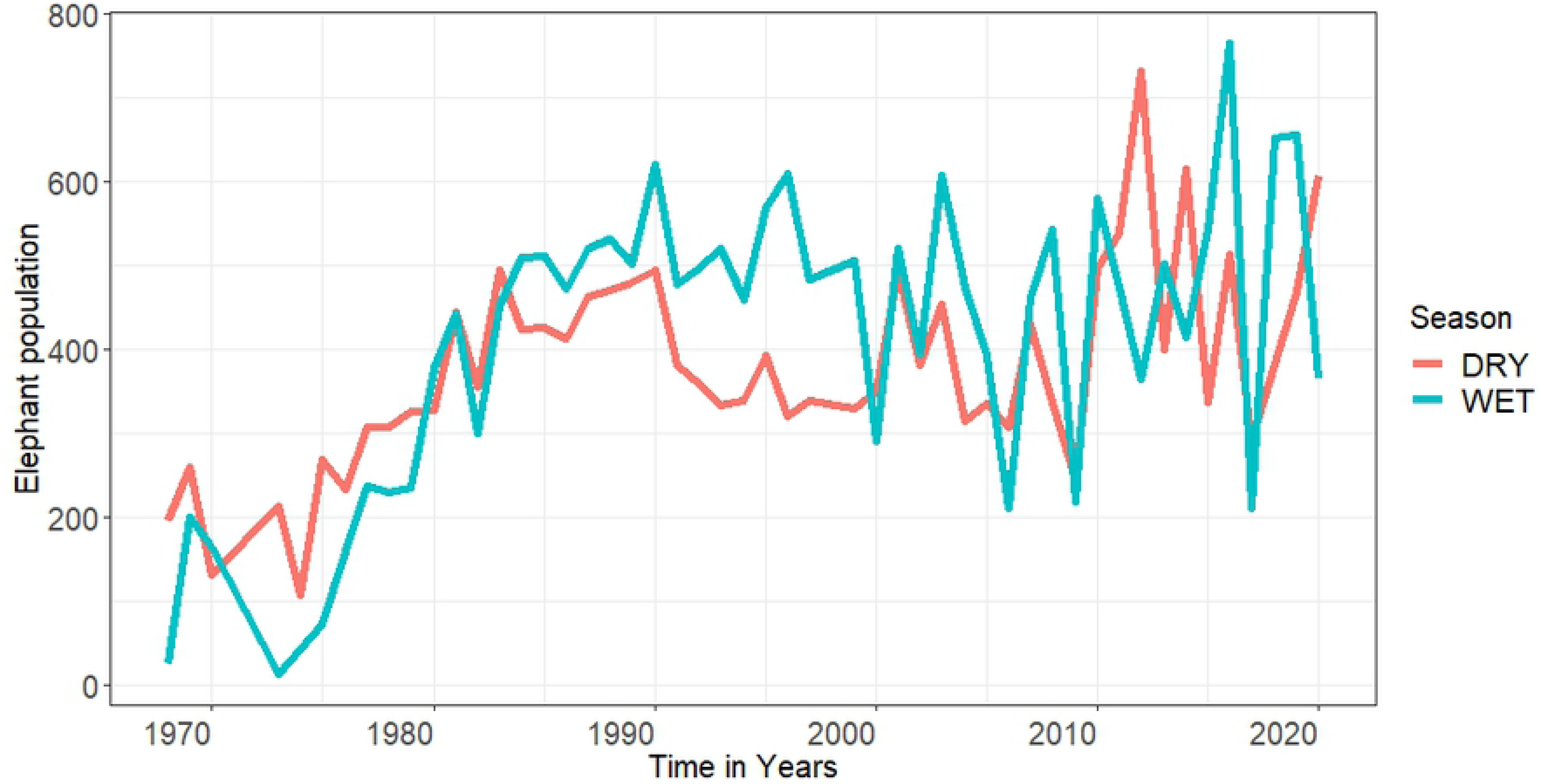
Mean annual wet and dry season population counts of the Amboseli Basin showing a drop in migrations in late 1970s and early 1980s in response to a surge in poaching outside the national park, followed by a reversal of migrations through to 2006 when movements became more erratic in response to wet and dry years.

Elephant protection in Amboseli National Park and surrounding group ranches spurred a recovery of the population from under 500 in the late 1970s to over 1,200 in the early 2000s. The recovery recorded by the aerial counts matches the growth in population to 1,500 in 2008 based on individually known animals documented by the Amboseli Elephant Program, many of which move beyond the ACP monitoring area (52). Despite the heavy losses of wildlife (54) and over 400 elephants in the drought of 2009 (Moss pers comm), the population continued to expand to nearly 1900 by 2020 (79), higher than the peak numbers recorded in the 1970s. Satellite tracking showed the enlarged population moving well into Tanzania and the Rift Valley (80), far further than radio-tracked elephants ranged in the 1970s (63).

### Changes in seasonal migrations

From the 1960s through the late 1970s elephants spread across the ecosystem during the rains and returned to the Amboseli Basin in the dry season, a regular seasonal migration also followed by wildebeest, zebra, buffalo and Maasai livestock (81). The seasonal movements track a plant quality gradient from the outlying bushlands during the rains to the woodlands of the Amboseli Basin in the early dry season and swamp-edge and swamp habitats in the late dry season (63, 82) as water sources and forage declined in the dispersal areas. The detailed feeding energetics of elephants studied by Lindsay (2011) shows a high rate of intake on taller grasses in the rains and a shift to woody vegetation and swamp sedges in the dry season. The seasonal movement of elephants and other migrants along the quality gradient has been explained by optimum foraging theory and allometric diet quality selection (84–86).

In a departure from the seasonal migrations which have continued unbroken by other species, elephant migrations halted abruptly with the heavy ivory poaching of the 1970s. The halt in wet season migrations caused a sharp rise in mean monthly population in the Amboseli National Park population (Fig 4). The change in seasonal migrations is captured in the monthly total counts of elephants in the Amboseli Basin. Between 1968 to 1978, prior to cessation of migrations, wet season counts of the Basin were significantly lower than rain season counts (*W* = 32, *P* = 0.0259). From 1980 to 1983 when the migrations halted, wet and dry season counts showed no significant difference (*W* = 4, *P* = 0.3429).

From 1984 to 2005 the migrations resumed but in reverse, with the dry season significantly lower than the wet season (*W* = 42, *P* < 0.0001). The seasonal differences became more erratic between 2006 and 2020 as peak numbers in the Basin oscillated between wet and dry seasons (Fig 4).

The population slump, cessation of migrations, compression of elephants into the national park and defensive herding behavior of elephants leaving the park all point to the disruptive impact of poaching in the 1970s. The compression was evident in the five-fold increase in elephant numbers in the park and the impact on the late season forage in the swamps. The growing forage shortage coincides with steady decline in elephant numbers in the park from an annual peak of 618 in 1990 to a low of 218 in 2009 during an extreme drought year when over 400 elephants died during the extreme drought (Moss, pers. comm), along with the majority of zebra, wildebeest and livestock (54). The decline in elephant numbers in the Amboseli Basin over the twenty-year period correlates with the decline in plant biomass in the basin (*r* = 0.78, *P* = 0.01). Elephant numbers in the park rebounded with the regeneration of pastures after the drought and only began to fall again during the dry years running into 2017. Numbers in the park grew once more between 2018 to 2020 when the swamp forage rebounded with heavy rainfall years.

### Compression along the seasonal foraging gradient

The concentration of elephants in Amboseli National Park in response to poaching created a knock-on effect along the seasonal habitat feeding gradient. The rising numbers initially built up in the woodlands, then shifted to the swamp habitats, creating a cascade of plant and animal changes. Fig 5 shows the seasonal movement from bushland through woodlands to swamp documented by Western and Lindsay (1984) intensifying with the surge in elephant numbers in the park during the 1970s and 1980s, rising five-fold in the swamps by 1995.

**Fig 5:**
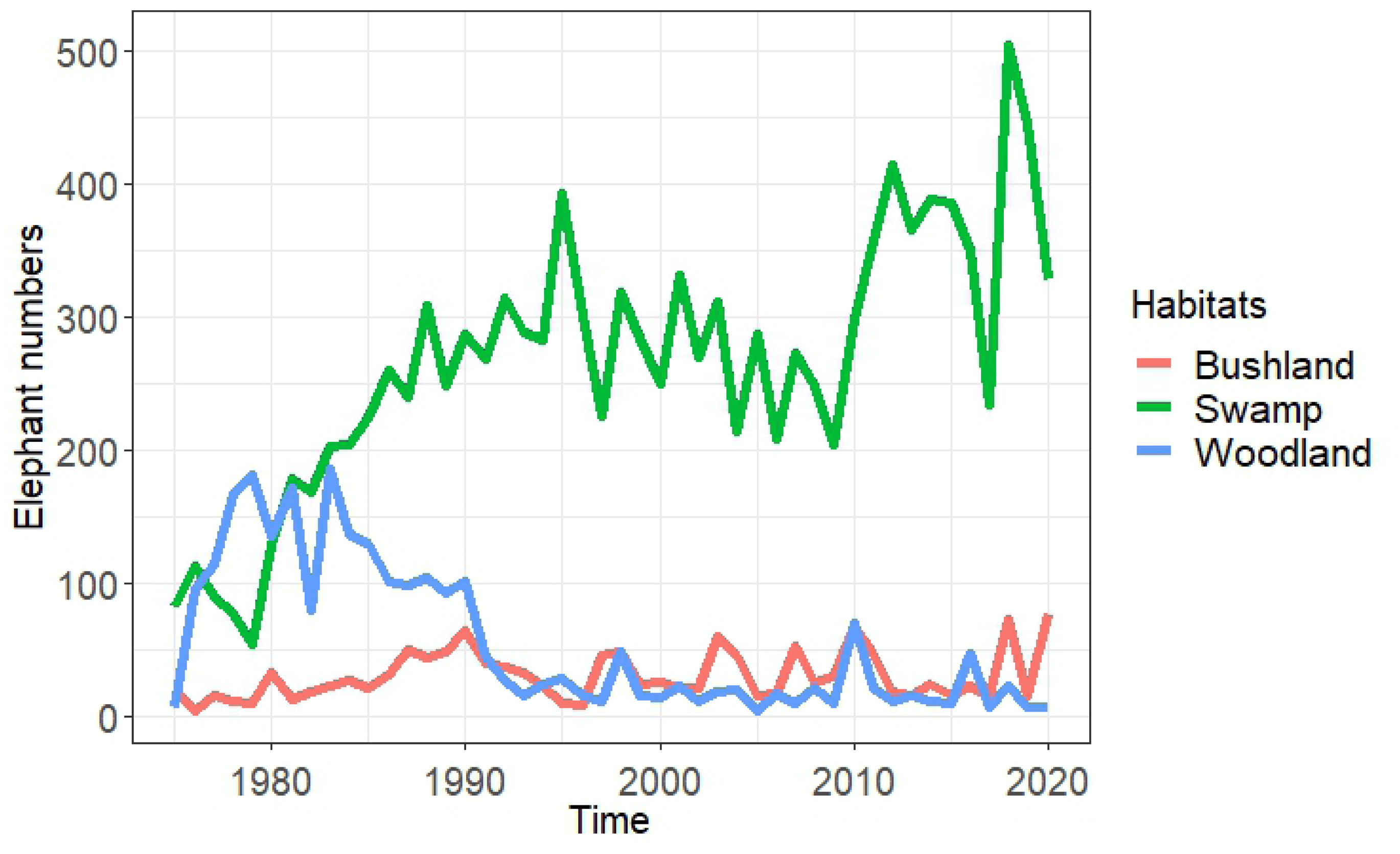
Elephant habitat use in response to poaching shifted progressively along the selectivity gradient from bushlands to woodland, intensifying the use of the late season swamp-edge habitat and permanent swamp habitats in the Amboseli Basin.

The loss of *Acacia xanthophloea* woodlands in the park saw a proliferation of low shrubs dominated by *Salvadora persica* and *Azima tetracantha* in the early 1970s. As the elephant numbers continued upward, *Salvadora* and *Azima* bushes were thinned and heavily pruned, causing the spread of more browse-tolerant species, including the saltbush, *Suaeda monoica,* small herbs dominated by *Dicliptera albicaulis,* and the tall coarse grass *Sporobolus consimilis.* The continued rise in elephant numbers led to growing browsing pressure on *Suaeda* and a gradual shrinkage in *Suaeda*-dominated shrublands in the early 1990s (87). In response to the loss of woody vegetation, elephant density in the swamps rose sharply (Fig 5). The shift from woodland to swamps correlates inversely with the depletion of total woody biomass in the Basin between 1975 and 1995 (*r* = −0.49, *P* = 0.01).

The heavy feeding and trampling of elephants in the swamps as mean monthly numbers rose from 83 to 392 over the twenty years from 1975 saw fringing woodlands and 2m tall thickets of *Solanum incanum, Pluchia dioschordes and Withania somnifera* herbaceous cover replaced by a dense sward of *Cynodon dactylon* and *Digitaria scalarum* grasses. The elephants also pushed deeper into the 2-3m tall *Cyperus papyrus, Cyperus immensus* and *Typha domingensis* sedges in the permanent swamps, resulting in the invasion of *C. laevaticus* and *C, merkii* and floating mats of herbs such as *Hydrocotyle ranunculoides*, *Ludwigia stolonifera* and the Nile cabbage, *Pistia stratoites* (88). By the early 1990s the intensive use of the swamps had opened up large stretches of open water and floating banks of vegetation. By the 2000 the heavy impact of elephants on the swamps caused an outward spread across the Amboseli Basin (Fig 6) and beyond (Fig 1), an outward spread also documented by the Amboseli Elephant Research Project (80).

**Fig 6:**
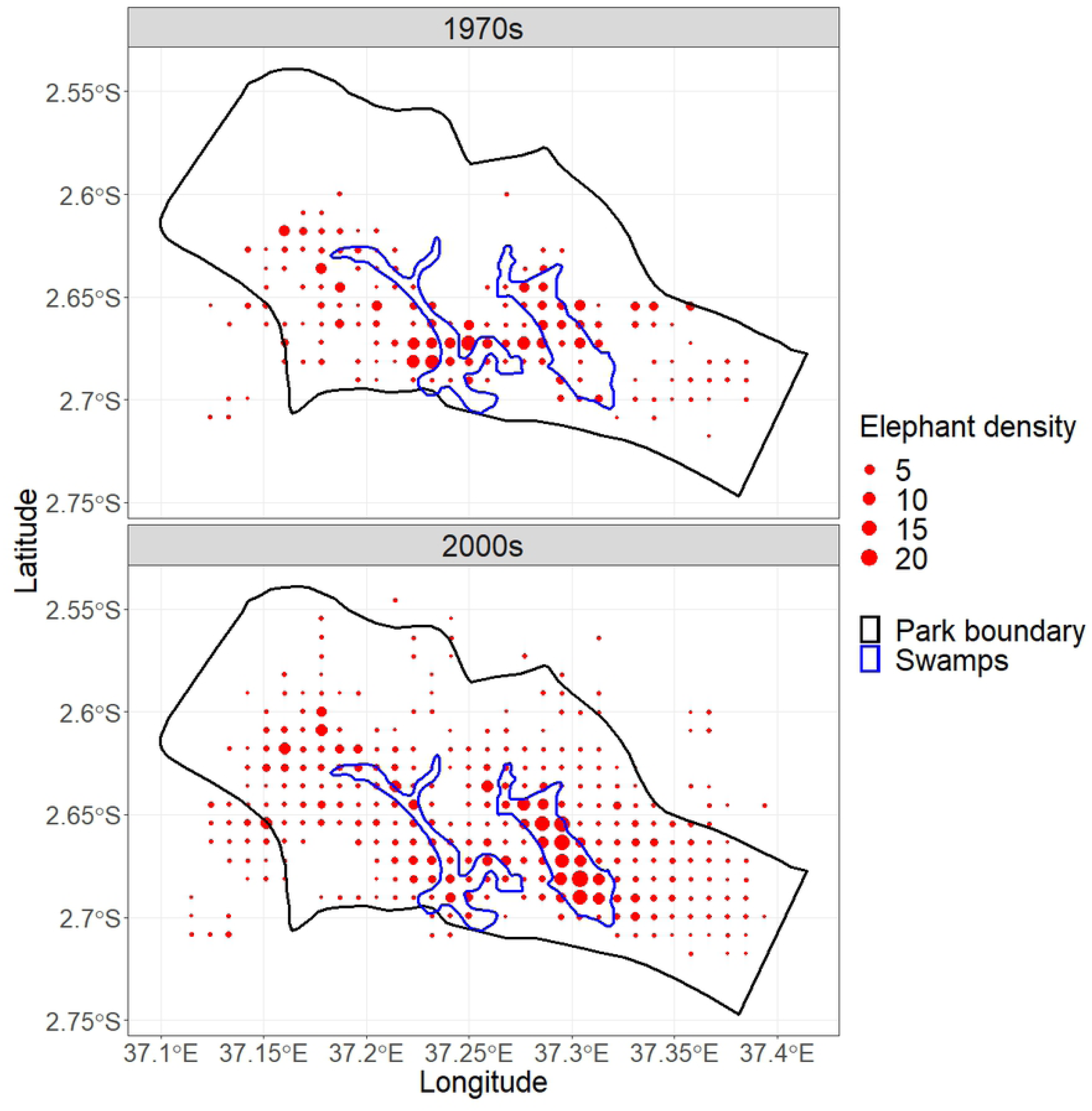
The compression of elephant into the swamp-edge and swamp habitats (Fig. 5) depleted pasture biomass, causing herds to forage more widely within and beyond (Fig. 4) the Amboseli Basin by the early 1990s.

### Elephant impact on the Amboseli woodlands

The loss of woodlands in Amboseli mirrors the rising elephant numbers from sporadic sightings in the early 1950s (47) to the large numbers presently. A study by Western and Van Praet (1973) showed elephants destroying *A. xanthophloea* by debarking mature trees, pushing over saplings and plucking seedlings. Although they attributed the cause of woodland loss to salinization accelerated by elephant damage at the time, later reanalysis of their data using partial correlation discounted salinization. A 20-year multifactor exclusion experiment showed elephants to be the primary cause of loss and the lack of regeneration (53).

The close correlation between elephant density, tree damage and tree death found by Western and Van Praet (1973) makes the fever tree a useful indicator species for tracking the ecological impact of elephants in the Amboseli Basin over the last seventy years using aerial photographic coverage. We selected photo plots on 1950, 1980 and Google Earth imagery (89) for 2020 to quantify the changes in woodlands density along transects running east and west from the Basin center at 1-kilometer intervals. The results (Fig 7) show fever woodlands in 1950 were dense and relatively uniform across the basin. By 1980 tree density had fallen steeply in the national park, corresponding to the sharp rising elephant numbers in the Basin (Fig 4). Tree survivorship increased with distance from the boundary and inversely proportional to elephant density (*r_s_* = 0.67, *P* = 0.0064). No trees remained standing in the park by 2020, except for two small groves, one in Ol Tukai Orok partially protected by the palm tree, *Phoenix reclinata*, the other bordering the Enkongo Narok swamp close to dense Maasai settlements (87). Despite the extinction of woodlands across most of the Basin over the seventy-year period, the distant-most groves outside the park remained vibrant.

**Fig 7:**
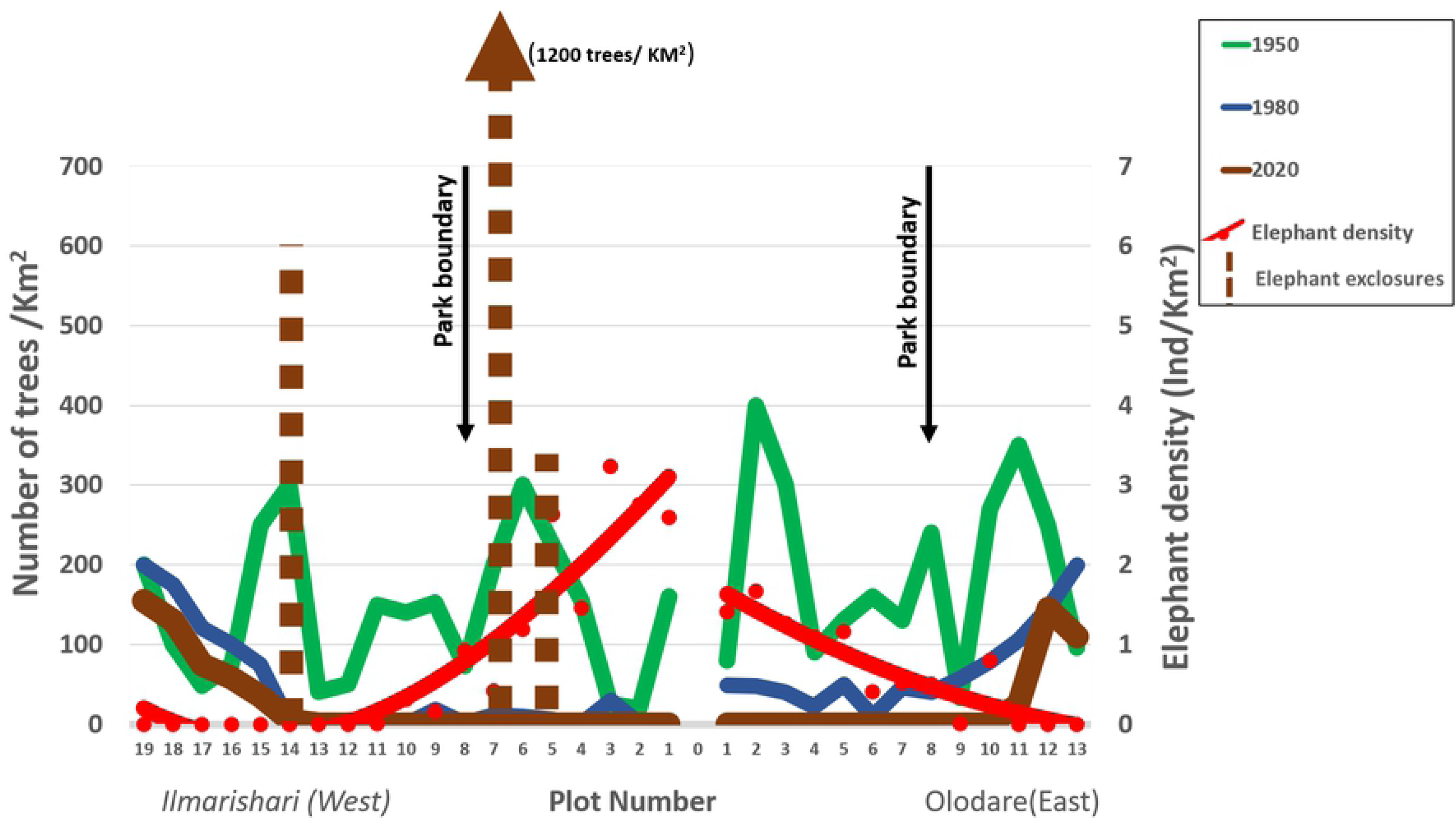
Density of fever trees in transects from center to periphery of Amboseli National Park from 1950 to 2020. Woodland density was uniform across the basin in the 1950s and declined steeply with the influx of elephants into the park in the 1970s. The pattern of woodland loss and regeneration reflects elephant impact which declined with distance from park boundary. Elephant-proof exclosures have reestablished dense regenerating groves.

A repeat measure correlation analysis shows that the declining woodland loss from the park center to basin periphery correlates highly with elephant density (*r_rm_* = −0.62, *P* = 0.0298), confirming the density-dependent tree damage found by Western and Van Praet (1973). The persistence of dense mature woodlands in the eastern periphery of the Basin among the irrigated farms, and in the western periphery where poaching and hunting across the border in Tanzania deterred elephants for several decades, points to the mitigation of their impact in woodlands where they are displaced by human activity.

High-level electric fences have shown how rapidly woodlands can be restored when elephants are excluded. A twenty-year experiment from 1981 and 1990 showed rapid tree recovery to maturity (S1 Fig) when elephants were excluded by a high-level electric fence allowing in meso-browsers (53). Later experimental exclosures showed similar results. These included a 0.4ha elephant exclosure built at Ilmarishari (S2 Fig) inside the park by ACP in 2001, two constructed by Kenya Wildlife Service (KWS) at Simek and Ol Tukai Orok in 2010, and two outside the park at Soito Nado by Ker and Downey Safaris between 2000 and 2010. All three elephant exclosures grew dense groves of fever trees within five years. The density of young regenerating trees in the ten-year KWS exclosures far exceeded the density in the mature 1950 groves. Trees in the ACP exclosure have matured, thinned and are approaching the density of the mature groves counted on the 1950 aerial photos. Fifteen other exclosure across basin established by KWS and tourist lodges since 1990 show strong tree regrowth against the continuing loss of woodlands and woody vegetation in the park.

The density-dependent tree damage creating an outward wave of woodland loss from the basin center to periphery (Fig 7), coupled with the recovery of woodlands in the elephant-proof exclosures, underscores the impact of elephants on the acacia woodlands. We next look at the wider impact on habitats and species composition.

### Changes in habitat and species composition

The most significant habitat changes in Amboseli since the 1950s have been described in detail in Western (2007) and Croze and Lindsay (2011). From 1950 to 2017 grassland habitats expanded from 28% to 40% of the Basin area in inverse proportion to the contraction of woodlands from 25% to 5% (*r* = −0.96, *P* < 0.0001). The fever tree woodlands were supplanted by a *Suaeda*-dominated shrubland. The heterogeneity in woodland structure and species composition converged over the period from the 1970s to 2017, due largely to the woody mass declining across the Basin from 600 gm^-2 to^ 200 gm^-2^ (72). The permanent swamps increased by two-fold, switching from tall to short sedges and banks of floating weed and large stretches of open water (88).

Both naturalistic and experimental exclusion experiments can be used to deduce the role of elephants in the changing habitats and species composition since the 1950s.

In the naturalistic experiment we used the 1988 combined ground transects running east and west from the park center to Basin periphery (Fig 7) to test whether grass and shrub biomass increased with declining elephant density along the transects. We found both the ratio of grass (*y* = −0.6*x* + 9.3, *R*^2^ = 0.54, *P* = 0.0061) and shrubs (*y*_1_ = −0.28*x* + 2.6, *R*^2^ = 0.49, *P* = 0.0118) to decline in proportion to tree biomass density (See Appendix A1)

In our controlled experimental manipulation of elephant impact, we used the high-wire exclosures in the former woodland areas to test whether elephant exclusion would revert the *Suaeda*-dominated shrublands and to fever tree woodlands. The ratio of both shrub (*y*_*s*_ = −0.9*x*_*e*_ +1795, *R*^2^ = 0.41, *P* = 0.0341) and grass biomass (*y*_*e*_ = −7.42*x*_*e*_ + 14881, *R*^2^ = 0.38, *P* = 0.0158) fell sharply with the increase in tree biomass (See Appendix A1). By 2015 the woodlands had formed a nearly closed canopy of mature trees. The shade-intolerant *Suaeda* bushes grew tall and thin in the woodland understory and, except for a few in the light gaps, disappeared from the maturing woodlands by 2015.

The highwire Ilmarishari electric fence, which also enclosed a swamp heavily grazed and trampled down as with other swamps in Amboseli, was used to test whether elephant exclusion would reestablish the tall-sedge swamps suppressed by heavy grazing and trampling as well as the fever tree woodlands. The enclosed swamp was monitored for changes in plant biomass and composition between September 2002 and July 2005 (88). An adjacent unfenced swamp 250 distance was monitored as a control. Despite no change in the control swamp, the residual *Cyperus papyrus* and *Cyperus immensus* sedges began growing back in the center of the experimental exclosure within a year, even as zebra and wildebeest maintained the short grazing lawns along the shallow swamp margins dominated by the grasses *Cynodon dactylon* and *Digitaria scalarum* (88). The exclosure showed a full recovery of 3m-tall sedges within five years of exclosure. Sakar (88) concluded that elephants play a major role in the ecological modification of wetlands as well as woodlands in Amboseli and more widely in Africa.

The changes in plants species composition and ecological function in the Amboseli Basin over the last five decades have been detailed by Western et al. (2021). The most significant changes have been a reduction in habitat diversity, convergence in species composition among habitats to smaller plants, and as a result, a decrease in species biomass, increased turnover rate and greater species dominance by herbivore-resilient species of shrubs and grasses. Plant production per unit of rainfall and resilience also declined significantly.

The overall changes in species richness, including trees, shrubs, herbs and graminoids in response to elephant density along the combined east and west transects across the Basin is shown in Fig 8. Species richness increased inversely proportional to elephant density from the park center to boundary (*r_s_* = 0.67, *P* = 0.0064), before declining sharply with increasing distances from the park boundary.

**Fig 8:**
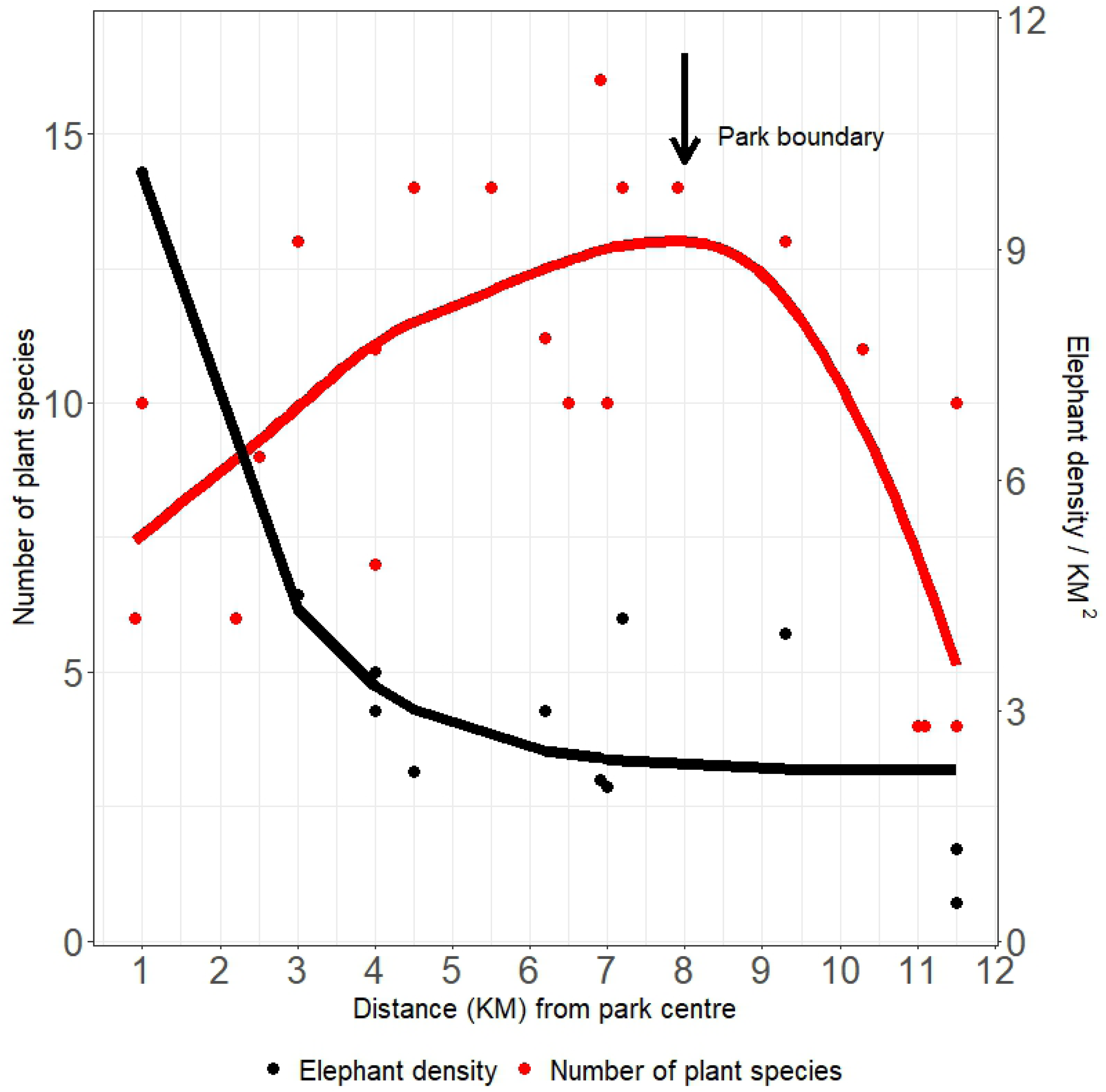
Plant species richness in relation to elephant densities along combined east and west transects from the center to periphery of the Amboseli Basin. Plant richness increases linearly with declining elephant density to the national park boundary then declines sharply at increasing distances.

### Herbivore ecological cascades

The changing composition of the large herbivore community in response to habitat changes in Amboseli has been tracked over the 40-year period from 1964 to 2004 using the surface bone assemblages which match the living population composition of 15 large herbivores species (66). The results showed a strong decline in the browsing relative to grazing community and the changes in guild composition to spatially track the transition from woodland to grasslands and expansion of the swamp grazing lawns.

The composition of the herbivore guilds in response to the habitat changes caused by intensified elephant use of the Amboseli Basin is captured by comparing ground counts in the 1960s and 2010s (Fig 9). In the 1960s elephants largely used the plains early dry season and woodlands in mid to late dry season. Elephants made little use of the swamps which were then dominated by tall sedges and a surrounding thicket of *Solanum incanum, Pluchia dioschordes and Withania somnifera* herbaceous cover. The seasonal shift of all herbivores through the habitats in the 1960s has been described in detail elsewhere (56). Browsers, including over 100 giraffe and 800 impala were regularly counted in the woodlands along with small numbers of bushbuck, lesser kudu and gerenuk. By the 2010s, following the loss of woodlands and spread of *Suaeda* bushlands, zebra, wildebeest and Thomson’s gazelle predominantly foraged on the swamp-edge grazing lawns created by elephants (Fig 9). Fewer than 10 giraffes entering the Basin for water are ordinarily recorded on current monthly counts. Impala numbers have fallen to fewer than 200, and bushbuck and lesser kudu have disappeared from the Basin.

**Fig 9:**
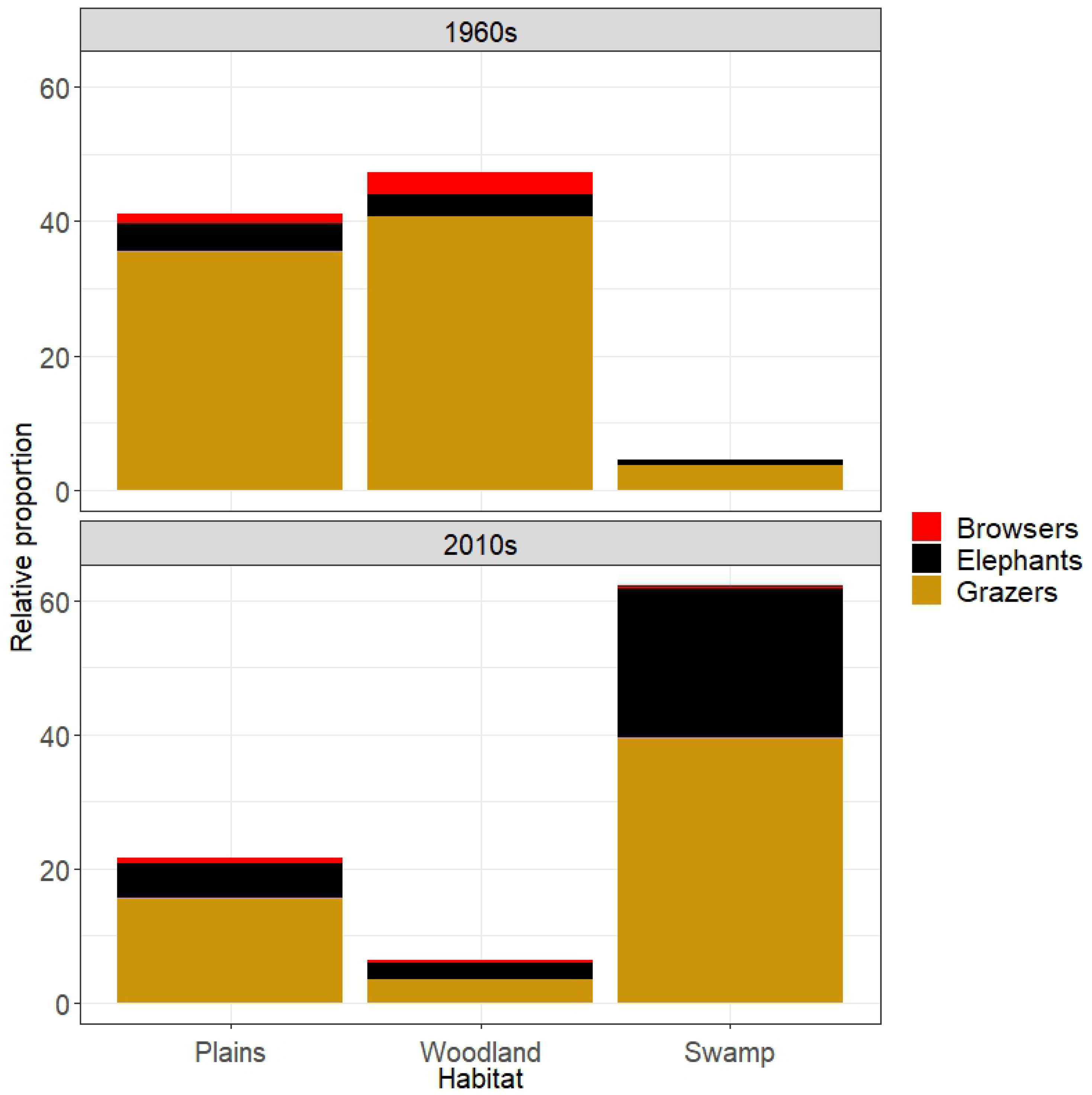
The loss of woodlands and spread of *Sueada* shrubland shifted elephant foraging to heavy year-round use the swamps, resulting in the tall sedges and shrubs giving way extensive grazing lawns. Browser numbers fell in response to woodland loss and grazers numbers increased as they moved onto the newly created swamp grazing lawns opened by elephants.

The shifting use of habitats by herbivores in response to the changes in the plant community over the past five decades can be analyzed by the use of a Detrended Correspondence Analysis (DCA) shown in Fig 10. The main changes show the grazing guild dominated by zebra and wildebeest to shift from a predominant use of the plains and woodlands to a heavy concentration in the swamp habitats in association with elephants. The browsing guild, including giraffe and Grant’s gazelle, shifted out of the basin to the bushlands. Rhinos a dominant feature of the browser guild were exterminated by poachers. Livestock use of the Basin habitats, including the central swamps, declined sharply following their exclusion from Amboseli National Park in the late 1970s.

**Fig 10:**
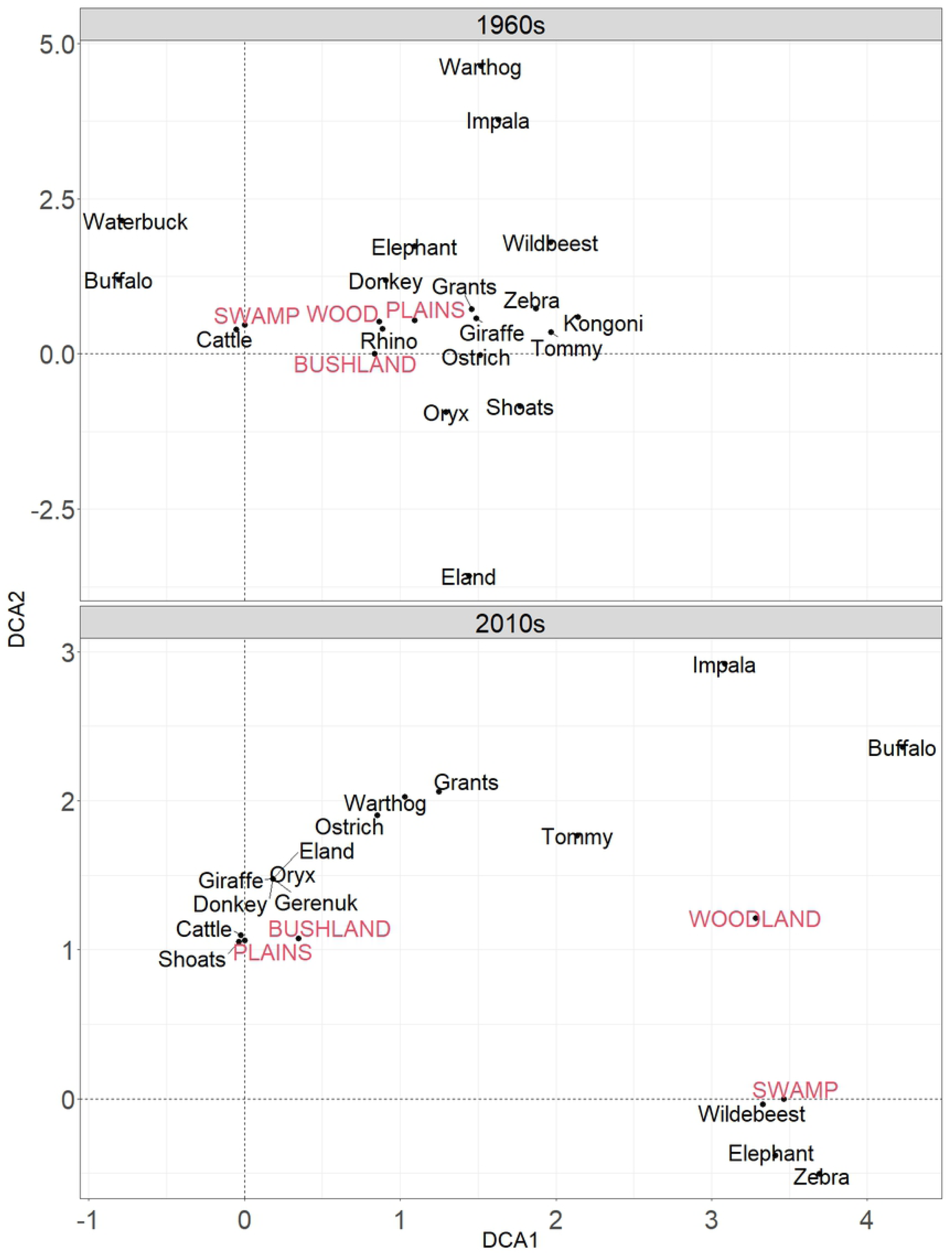
Detrended Correspondence Analysis (DCA) for the Amboseli large herbivores in the 1960s and 2010s. The swamps, which in the 1960s were used mostly by waterbuck, cattle and buffalo, are now dominated by elephant, wildebeest and zebra due to the transformation of the swamps from tall sedges and herbaceous cover to grazing lawns dominated the *Cynodon dactylon*. Browsing species have shifted to the bushlands surrounding Amboseli along with livestock which have been excluded from the Basin due to the establishment of Amboseli National Park.

The changes in herbivore species composition and ecological function in the Amboseli Basin over the last five decades have been detailed by Western and Mose (2021). Species diversity declined with the increase in grazers and decrease in browsers, causing large scale changes in the macroecology of the Amboseli herbivore community. Further, despite the increase in overall wildlife production due to the expansion of swamp grazing lawns, the probability of extreme forage shortfalls and herbivore population collapses increased due to the heavy sustained browsing and grazing pressure depressing plant resilience to extreme events (90). The macroecological changes reflect the increasing dominance of elephants which rose from 8% of total herbivore biomass in the Amboseli Basin in the 1960s to 29%, causing a cascade of changes in the plant and animal community.

## Discussion

The compressed and growing elephant population in Amboseli National Park has transformed woodlands into shrublands and expanded the area of grasslands and swamps over the last seven decades. The collapse of the migrations and compression of elephants into the park caused a cascade of floristic changes, including a convergence in composition typified by smaller browse-tolerant species, reduced primary production, an accelerated turnover rate and reduced resilience. Changes in the herbivore community included a decline in browsers associated with the woodland to grassland transformation.

Changes in the swamp vegetation highlight the keystone role of elephants as ecological engineers in opening up wetlands to a range of smaller herbivores. Sarkar (2006) has documented the effects of elephant trampling, fecal deposits, water aeration and sediment churning on swamp vegetation. The intensive herbivory boosted the growth rate and quality of the grazing lawn pastures, explaining the attraction of the swamps in the wet seasons between the 1980s and 2000s (Fig 4) when elephants would otherwise have migrated to the bushlands (83). The post 1980s decline in elephant numbers in the park during the dry season when elephants previously congregated in the swamps is explained by the depletion of forage due to heavy grazing (54), culminating in the heavy elephant and ungulate mortality in the 2009 drought (90).

We attribute the expansion of the swamps to the reduced transpiration rates and increased puddling caused by heavy grazing and trampling of the tall sedges, by flash floods into the Basin caused by sedentarization and heavily grazing in the surrounding group ranches (69), and perhaps to increased discharge of aquifers into the Basin from Kilimanjaro (52).

The impact of elephants on the Amboseli swamps mirrors Vesey-FitzGerald (1960) account of the grazing succession in Lake Rukwa in Southern Tanzania. Here, elephants trample and graze down tall sedges, creating a succession of smaller herbivores onto the newly created grazing lawns as the dry season progresses. The Amboseli findings also match reports of high-density elephant populations in protected areas. Guldemond and Van Aarde (2008) in a review of 238 studies and a meta-analysis of 21 research sites across Africa found high densities, amplified by low rainfall and fencing, to reduce woody vegetation.

The Amboseli points to a more complicated picture than compressed populations studies when elephants are free to move in response to humans at an ecosystem and larger scale. Transects across the Amboseli park boundary show a loss of plant species richness at low as well as high densities due to dense woodland cover suppressing understory plants. Poulsen et al. (2017) also found elephant extirpation to result in plant species loss and ecosystem simplification in African forests. Based on the density-dependent responses of plant species richness, we deduce that protected area research and compressed populations mask the larger shifting patchwork effect of elephants as landscape agents (94). The impact of compression and curtailed movements on woody vegetation is analogous ecologically to the impact of heavy sustained grazing on grassland production in response to land subdivision and sedentarization of pastoral herds on the Kaputei group ranches north of Amboseli (69, 72)

Although Guldemond et al. (2017) and Cook and Henley (2019) found high-density elephant populations do generally cause tree loss and habitat simplification, the ecological cascade effects can also be positive as in the case of the grazing success in wetlands. Skarpe and Ringrose (2014), in a detailed multispecies study of Chobe National Park in Botswana, added to the complex view in showing elephants to cause a wide range of cascade effects as a function of distance from water. Heavy localized impact of elephants on vegetation near water courses and waterholes caused an expansion of short grasslands, increase in meso-herbivores, large carnivores, small mammals and gallinaceous birds. Further from water lower elephant densities resulted in heavy bush and woodlands cover. The study also found elephant impact to be moderated by bottom-up geomorphology, soils and hydrology.

Fritz et al. (2002) in investigating the impact of megaherbivores on the guild structure across 31 African ecosystems found elephants to compete with meso-browsers and mixed feeders but not grazers, a finding echoed in our study. In a yet broader review of the role of megaherbivores, Bakker et al. (2016) combined paleo-data and exclosure studies to show that megaherbivore extinctions and exclusions can cause large ecological cascade no less than high-density populations, in this case through an expansion of woodlands and increase in species dominance.

The focus on protected areas and a lack of studies on free-ranging populations interacting with humans at a landscape scale has fostered a view of park elephants as being incompatible with biodiversity conservation. Elephant reduction programs have been used to alleviate the biotic impact. It is worth noting that many studies on the impact of elephants on vegetation were conducted during the 1970s and 1980s when herds were heavily poached and retreating to the safety of parks, as in Amboseli. The findings point out the inadequacy of protected areas in conserving biodiversity due to the compression of elephants and lack of large-scale movements.

In contrast, Hoare and Du Toit (1999) found elephants and humans coexisted across a wide-range of settings below a threshold of disturbance. They note the little attention paid to the ecological interaction of elephants and people moving freely in response to each other. Such evidence, and the findings from Amboseli, suggest the need for context-specific studies to decipher the ecological role of unrestricted elephant populations.

Elephants, we suggest need far larger space and the creative tension with human disturbances lacking in parks to play a positive keystone role in creating the patchwork of habitats and successional stages creating the mosaic of habitats supporting biodiversity in the African savannas and forests.

Contemporary elephant populations tell us even less about the ecological role of free-ranging elephants prior to colonial parks and the rapid human population growth in Africa over the last century (28, 98). Laws suggests elephants in pre-colonial days shifted with the loci of human activity, creating a large-scale mosaic of habitats with local differentiation due to browsing and grazing and human activity. Based on a multivariate 20-year elephant exclusion study in Amboseli, Western and Maitumo (2004) similarly suggested that elephants and livestock moving freely around the landscape set up a creative tension causing a shifting patchwork of habitats.

The creative tension is captured by the continuous “jostling” of livestock and elephants at the park boundary where species richness peaks due to the patchy habitats caused by a decadal ecological minuet. At the interface, elephants thin woodlands and livestock reduce grass and encourage woody invasion (53). The hump-back curve of plant species richness along elephant gradient (Fig 8) reflects the interplay of safety and threat responses of elephants to humans. The interplay explains the hump-back curve of plant species across the park boundary (Fig 6) where human-moderated elephant and livestock movements create a shifting tapestry of patchy habitats.

The species richness along the Amboseli elephant gradient fits the Intermediate Disturbance Hypothesis (99), as modified by the Milchunas-Sala-Lauenroth (MSL) models (100) along a productivity gradient. Although the literature shows elephants depleting woodlands and creating browse traps of dwarfed woody vegetation in high density park populations (101), the impact of short-term intensive browsing has yet to be studied at an ecosystem scale.

Our observations in Amboseli suggest that short intense browsing due to seasonal elephant movements and in response to human activity may stimulate woody production and diversity in much the same way that short-term intensive grazing can create browsing arenas which stimulate grass production (102–104) and species richness (100). Our observations of giraffe browsing lend support to this view. Heavy giraffe browsing on young fever tree stands creates a dense thicket of thorns which prevent elephants penetrating the barrier and debarking and trees until they reach near maturity, the apical branches reach above giraffe height and the lower densely thorned branches die back.

The context-specific impact of elephants points to the futility of separating the ecological role of elephants from people, treating parks as natural ecological systems and ignoring the importance of mobility and scale in ecological cascades (7, 8). The million years or more of human-elephant coevolution has shaped the ecological, behavioral and cultural adaptations in elephants and people. Elephants readily distinguish human responses by, for example, foraging beyond park boundaries at night to avoid people (23), yet acting benignly around safari vehicles, lodges and campsites in parks.

The response of people to elephants similarly varies, most notably in the cultural traditions and familiarity with elephant behavior. In Amboseli Maasai pastoral practices and tolerance of wildlife has fostered coexistence (105), whereas Maasai who have taken up farming and suffer crop raiding have become fearful and intolerant of elephants (106). In the same vein, human activity globally, whether promoting ivory trading for profit, or elephant protection and viewing arising out of modern conservation sensibilities, have become the predominant forces shaping the ecological cascades.

Other factors bearing on how elephants and people shape ecosystems include physical factors such as rainfall (20), landscape heterogeneity (107), the size of parks, distance from water (96) and seasonal migrations. Rainfall seasonality as much as total rainfall may also influence the impact of elephants on vegetation in eastern Africa. Here the bimodal rainfall supports shorter sparse bushlands and scattered grasslands in contrast to the taller denser deciduous woodlands of southern Africa’s unimodal rainfall regime.

The loss of elephant mobility and range and the rising conflict has reduced the scope for sharing land with people. The loss of coexistence has shifted the interaction from a positive perturbation promoting heterogeneity to ecological disruption and uniformity. Caughley (1988) in reviewing the causes and consequences of local overabundance in mammals suggested the two exceptions to the reversion of vegetation from temporary overabundance are due to the loss of mobility in livestock and elephants. O’Connor et al. (2007) goes further in suggesting elephants to be predominantly grazers based on their digestive physiology but increasingly browsers under range compression in protected areas.

We propose that prior to the ivory trade the creative tension of free-ranging elephants in response to humans created a patchwork of habitats and a shifting mosaic of vegetation consistent with the view of elephants as a keystone species and ecosystem engineers. The impact of the ivory trade over the last few centuries (37, 38), a breakdown in traditional lifestyles and cultural practices in land husbandry which sustained biodiversity over the last 12,000 years until industrial times (109), and the growth of protected areas, has subsequently compressed elephant ranges and caused a growing ecological dislocation by disrupting the keystone species (110).

Tempering the ecological dislocations of mega-mammals caused by local overabundance (111), reviving their keystone role in sustaining biodiversity (112), and reducing human-wildlife conflict impeding ecological upscaling (18, 113) can be achieved naturally by ensuring elephants and other large herbivores and carnivores have the space needed to maintain large self-sustaining populations. This is a formidable challenge though by no means insurmountable.

The widening horizons of conservation in the last few decades have seen protected and wildlife management areas win space for large mammals through community-based conservation initiatives (114, 115). Cumming et al. (2013) note that matching the scale of human demands on an ecosystem and the services the ecosystem can deliver calls for appropriate institutional frameworks for the scale-matching.

Examples of scaling up from park to ecosystem and the expanded institutional frameworks include the cross-border Kruger link between South Africa and Mozambique (117), the greater Amboseli ecosystem in Kenya (118) and the Greater Yellowstone Coalition in the U.S. (119). Yet wider regional linkages include the Paseo Pantera Central-South America landscape (120), the Yellowstone to Yukon landscape across the U.S.-Canadian border Chester, 2015), and the four-borders Kavanga-Zambezi (KAZA) landscape in southern Africa (121). The theoretical framework for widening to regional and continental levels in the face of human impact and climate change, have been highlighted by Soulé and Terborgh (1999), Allen and Singh (2016) and Curtin (2015).

Such examples hold out hope of finding the space and the mobility elephants and other large herbivores and carnivores need to play vital keystone roles in sustaining biodiversity in an increasing human-dominated world, in alleviating the ecological disruption of compressed populations in parks, and in minimizing the need for intense species population management.

## Appendix A1

The ratio of grass to tree biomass density (*y*) reduced by 0.6 for every additional kilometer (*x*) away from the centre of the park, (*y* = −0.6*x* + 9.3, *R*^2^ = 0.54, *P* = 0.0061). For shrubs, the ratio (*y*_1_) reduced by 0.28 with a unit increase in distance (*y*_1_ = −0.28*x* + 2.6, *R*^2^ = 0.49, *P* = 0.0118).

In the controlled experimental manipulation of elephant impact, the ratio of grass to tree biomass density (*y*_*e*_) reduced by 7.42 over the years (*x*_*e*_), (*y*_*e*_ = −7.42*x*_*e*_ +14881, *R*^2^ = 0.38, *P* = 0.0158). That of shrubs (*y*_*s*_) reduced by 0.9 (*y*_*s*_ = −0.9*x*_*e*_ +1795, *R*^2^ = 0.41, *P* = 0.0341).

## Acknowledgments

We wish to thank the many wardens and the staff of Kenya Wildlife Service for their support of the Amboseli Conservation Program over the years. David Maitumo assisted in data collection for the experimental exclosures and plant monitoring. Eric Ochwangi, Rebecca Kariuki and Caroline Mburu helped analyze elements of the long-term data set we draw on in this paper. Winfridah Kemunto has compiled much of the field data for computer analysis. We thank Shirley Strum for her review of the manuscript.

## Author Contributions

David Western has designed, planned and directed the field programs of the Amboseli Conservation Program since its inception in 1967. Victor N. Mose has been responsible for data management, statistical analysis and graphical presentation the data used in this paper. Both authors contributed to the conceptualization of the paper and divided writing assignments. Western was responsible for the literature review and final manuscript writing, Mose for the preparation for publication.

## Data Availability Statement

The extensive data used in this paper was collected and is retained by the Amboseli Conservation Program at the African Conservation Centre, Nairobi, and can be requested via www.amboseliconservation.org or through the director and Database Administrator of the Amboseli Conservation Program at (|).

## Ethics statement

Kenya Wildlife Service (KWS) issued permits to conduct field work inside the protected Amboseli National Park (latitude: −2.626610; longitude: 37.2544060). For study sites located in the surrounding group ranches (Fig 1), no endangered plant species were involved. Aerial sample surveys of large herbivores were conducted in line with the Kenya Government, Department of Resource Surveys and Remote Sensing (DRSRS) methodology.

## Supporting information captions

**S1 Fig:** Exclusion trench and high-wire fence established in 1981 to test the impact of elephant removal on fever tree regeneration (A). The fever trees had grown to a mature woodland by 2003 (B) and the high-wire fence expanded in 1992 to include the lodges in the background.

**S2 Fig:** A high-wire electric fence set up in 2001 at Ilmarishari exclosure (A) to test the impact of elephant removal on woodland recovery. By 2008 the regenerating fever trees had shaded out the Suaeda-dominated shrublands which supplanted the woodlands destroyed by elephants in the 1970s. The aerial view of the exclosure in 2005 (B) showing extensive fever tree regeneration and regrowth in the swamps relative to the control open-water in the foreground. By 2018 the fever trees formed a dense canopy closing out *Suaeda* shrub (C).

